# Aldehyde dehydrogenase 3 is an expanded gene family with adaptive roles in chickpea

**DOI:** 10.1101/2019.12.17.879213

**Authors:** Rocío Carmona-Molero, Jose C. Jimenez-Lopez, Juan Gil, Teresa Millán, Jose V. Die

## Abstract

Legumes play an important role in ensuring food security, improving nutrition and enhancing ecosystem resilience. Chickpea is globally an important grain legume adapted to semi-arid regions under rain-fed conditions. A growing body of research shows that aldehyde dehydrogenases (ALDHs) is a gene class with promising potential for plant adaptation improvement. Aldehyde dehydrogenases constitute a superfamily of proteins with important functions as ‘aldehyde scavengers’ by detoxifying aldehydes molecules and thus playing important roles in stress responses. We performed a comprehensive study of the ALDH superfamily in the chickpea genome and identified 27 unique ALDH loci. Most chickpea ALDHs originated from duplication events and ALDH3 gene family was noticeably expanded. Based on the physical locations of genes and sequence similarities, our results suggest that segmental duplication has been a major driving force in the expansion of the ALDH family. Supported by expression data, the findings of this study offer new potential target genes for improving stress tolerance in chickpea that will be useful for the breeding program.

## 1. Introduction

Aldehyde molecules are common intermediates of a number of catabolic and biosynthetic pathways that are produced in response to biotic and abiotic environmental stresses. Although aldehydes are indispensable to developmental and growth processes, excessive amounts of aldehydes interfere with metabolism becoming toxic, so their unbalanced levels must be regulated within the cells (Bartels, 2001; Jakoby and Ziegler, 1990). The aldehyde dehydrogenase (ALDH) superfamily is a group of NAD(P)+-dependent enzymes that catalyze the irreversible oxidation of a wide range of reactive aldehydes to their corresponding carboxylic acids (Lindahl, 1992; Yoshiba et al., 1997). In addition, under conditions inducing oxidative stress, ALDH enzymes act as ‘aldehyde scavengers’ by metabolizing reactive aldehydes derived as lipid peroxidation-derived aldehydes, which are potentially toxic due to their extreme reactivity with the nucleophilic compounds such as nucleic acids, proteins and membrane lipids (Rodrigues et al., 2006; Yoshida et al., 1998). However, ALDH activity may also serve to fine-tune gene activation since ALDHs may modulate signalling by lipid peroxidation-derived bioactive aldehydes (Missihoun and Kotchoni, 2018).

Interestingly, ALDHs are found throughout all taxa including both prokaryotes and eukaryotes, where many ALDH families are highly conserved among animals and plants (Brocker et al., 2013). To date, ALDHs have been identified and categorized into 24 separate families based on protein sequence identity as main criteria (Black and Vasiliou, 2009), but also by their functionality (Jimenez-Lopez, 2016; Jimenez-Lopez et al., 2016). Since the first identified plant ALDH gene *rf2*, which encodes a mitochondrial class-2 ALDH, was reported to function as male fertility restorer (RF) protein of maize (Skibbe et al., 2002), and many others ALDH were classified as RF afterward (Kotchoni et al., 2010), a number of studies have demonstrated that ALDH genes are involved in diverse pathways and appear to play crucial roles in molecular detoxification, as well as growth and development (Guo et al., 2017; Kotchoni et al., 2012; Shin et al., 2009). In addition, many of the plant ALDH genes characterized to date are induced under wide range of abiotic stresses such as drought, cold, high salinity, and heavy metals highlighting their potential role in improving stress tolerance/ environmental adaptation (Bartels, 2001; Gao and Han, 2009; Kirch et al., 2005; Xu et al., 2013).

The identification of ALDH genes in different crop species has soared in recent times due to the increasing numbers of plant species that have been sequenced. Among the plant species containing 14 distinct ALDH families, the ALDH11, 12, 19, 21, 22, 23, and 24 are distinguished unique in the Plantae kingdom. The single gene of the ALDH19 family reported so far, encodes a gamma-glutamyl phosphate reductase involved in proline biosynthesis (García-Ríos et al., 1997); and no other higher plant has been found containing this family.

Chickpea (*Cicer arietinum* L.) is globally the second most important grain legume (FAOSTAT, 2018). Although the yield potential has increased over the last years, the global production is constrained by several major abiotic (drought, heat, high salinity) and biotic stressors such as the fungal diseases *Fusarium* wilt, and *Ascochyta* blight, which may cause 100% loss in yield when conditions are favourable for infection (Li et al., 2015; Millán et al., 2015). Until recently, lack of information on legume genomes traditionally restricted the survey of gene functionalities in response to the environment or stress, which may be valuable to be implemented in breeding programs for chickpea yield improvement under climate change immediate adaptation. Fortunately, the genome sequence of chickpea has become available in the last few years providing an unprecedented resource that can be exploited in numerous ways (Jain et al., 2013; Varshney et al., 2013).

In the present study, we identified 27 ALDH loci in the chickpea genome encoding a total number of 45 proteins that contained the complete ALDH domain and belonged to 10 different ALDH families. We performed a comprehensive functional comparison of the chickpea ALDH superfamily to other sequenced plant species, through phylogenetic and synteny analyses, the study of their expression profiles in response to various types of stress, and structure-functional analysis. Results from this study provide functional targets with yield improvement potential for the chickpea breeding program, as well as the basis for further comparative genomic analysis and a framework to study the ALDH genes evolution on a large timescale within the legume family.

## 2. Materials and methods

### 2.1. Database searches and annotation of ALDH genes

Comprehensive identification of *C. arietinum* ALDH gene family members was achieved using A*rabidopsis thaliana, Glycine max*, and *Medicago truncatula* ALDH proteins. A keyword-based search was carried out against the databases of the National Center for Biotechnology Information (NCBI) to extract 136 *A. thaliana*, and 55 *G. max* ALDHs. In addition, 36 *M. truncatula* ALDHs were downloaded from the Phytozome v12.1 database (https://phytozome.jgi.doe.gov). All these sequences were used as queries in BLASTP searches (Altschul et al., 1990) to identify the corresponding ALDH members in the chickpea proteome using a cut-off of query coverage ≥ 25%, E-value ≥ 1e-25, and identity ≥ 25%. The Pfam domain PF00171 (ALDH family), PS00070 (ALDH cysteine active site), PS00687 (ALDH glutamic acid active site), and the accession ‘cl11961’ were queried against the Pfam (https://pfam.xfam.org/), and the CDD (https://www.ncbi.nlm.nih.gov/cdd/) databases to confirm the candidate sequences as ALDH proteins. For exhaustive identification of divergent members, we used the chickpea sequences as queries in BLASTP searches against the chickpea proteome. These steps enabled us to obtain 45 unique ALDH protein sequences. Using one gene model per locus, we identified 27 *C. arietinum* non-redundant ALDH genes (CaALDH). Information on chromosomal location, locus ID, amino acid length, molecular weight, and number of exons was retrieved from the NCBI using the *refseqR* package (Die, 2018). The ExPASy proteomics server database (https://www.expasy.org/) was used to predict the theoretical isoelectric point (pI) of each ALDH protein, as well as the molecular weights (MW) of the deduced proteins without that record in the NCBI. For subcellular localization predictions and active site assessment, we used DeepLoc 1.0 (Almagro Armenteros et al., 2017), SLP-Local (Matsuda et al., 2005), SMART, ChloroP 1.1 (Emanuelsson et al., 1999), Mitoprot (Claros and Vincens, 1996), PROSITE and PROPSEARCH databases (Hobohm and Sander, 1995). Putative ALDHs were further annotated on the basis of the ALDH Gene Nomenclature Committee (AGNC) annotation criteria (Vasiliou et al., 1999). Briefly, amino acid sequences that shared > 40% identity to previously identified ALDH sequences were considered to comprise a family; those exhibiting > 60% identity comprise a protein subfamily, while sequences with < 40% identity are considered to be a new family.

### 2.2. Syntenic blocks and gene duplication analysis

Syntenic blocks between chickpea and *Medicago truncatula* genomes, were downloaded from the Plant Genome Duplication Database (Lee et al., 2013). Those containing CaALDH genes were identified and analyzed. Duplicated genes were labelled as ‘duplicated genes’ according to the criteria defined by (Tan and Wu, 2012): (1) the alignment covered >70% of the longer gene; (2) the aligned region had an identity of >70%. Coverage and identity values were obtained by BLAST searches of all the predicted CDS against each other. Tandem duplicated genes were defined as those closely related in the same family and clustered together within a sliding window size < 250 kb (Ameline-Torregrosa et al., 2008). The Circoletto tool was used to plot sequence similarity (Darzentas, 2010).

### 2.3. Phylogenetic analysis of ALDH gene families

To carry out the phylogenetic analysis, the alignments of the deduced amino acid ALDH protein sequences from *Arabidopsis* (Kirch et al., 2004), soybean (Kotchoni et al., 2012), maize (Jimenez-Lopez et al., 2010) and chickpea were performed using the ClustalW program as implemented in the Molecular Evolutionary Genetics Analysis software (MEGA) version 6 with default options (Tamura et al., 2013). The alignments were created using the Gonnet protein weight matrix. Sequences < 250 aa were eliminated from the matrix because short sequences interfered with a fine alignment. A total of 79 proteins were finally used to build the ALDH phylogeny of chickpea. The phylogenetic tree was constructed using the Neighbor-Joining method implemented in MEGA and the reliability of the interior nodes was assessed using 500 bootstrap replicates.

### 2.4. *In silico* expression analysis

The coding sequences of ALDH genes were used to query the NCBI chickpea ESTs. Searching parameters were set as follows: blast algorithm megablast, identity > 95%, query coverage > 25% and E-values < 10^−20^. Next, the full-length CDS of the ALDH genes were employed to query the NCBI Sequence Read Archive (SRA) database (https://www.ncbi.nlm.nih.gov/sra). For assessment of ALDHs expression support in response to the fungus *Fusarium oxysporum*, we selected two libraries constructed from infected root samples of resistant (WR315) chickpea plants after 48 hours post-inoculation (SRX535351), and control samples of resistant (WR315) chickpea plants (SRX535349) using Magic-BLAST as the mapper (Boratyn et al., 2019). The searching parameters were implemented as follows: only one read per hit was counted, length reads were equivalent to 100 bp, and the identity > 99%. Normalized counts of hits were performed using public scripts to quantify the expression of transcripts from datasets (https://github.com/NCBI-Hackathons/SimpleGeneExpression).

### 2.5. ALDH proteins modelling and structural features analysis

The ALDH protein sequences NCBI: XP_0045024851, XP_004502482, and XP_004498346 were used for searching the best structural templates in the Protein Data Bank (PDB). The suitable templates for these sequences were selected by BLAST server (http://ncbi.nlm.nih.gov/). We used the software BioInfoBank Metaserver (http://meta.bioinfo.pl/) to improve the best final templates selection, as well as the Swiss-model server for template identification (swissmodel.expasy.org). The best four templates were retrieved from PDB database 1ad3 (Liu et al., 1997), 4qgk (Keller et al., 2014), 5nno (Zhang et al., 2018), and 5ucd (Zahniser et al., 2017), and implemented for homology modelling. ALDH protein models were built by the implementation of SWISS-MODEL via the ExPASy web server (swissmodel.expasy.org). 3D structural errors recognition was performed by ProSA (prosa.services.came.sbg.ac.at/prosa.php), as well as first overall quality estimation of each model by QMEAN4 (swissmodel.expasy.org/qmean/cgi/index.cgi). Final ALDH structures were subjected to energy minimization using GROMOS96 and implemented in DeepView/ Swiss-PDBViewer v3.7 (spdbv.vital-it.ch) to improve the Van der Waals contacts and correct the stereochemistry. Final model’s quality was assessed by testing proteins stereology with PROCHECK (www.ebi.ac.uk/thornton-srv/software/PROCHECK), and ProSA programs, as well as the protein energy with ANOLEA (protein.bio.puc.cl/cardex/servers/anolea). Next, the Ramachandran plot statistics for the models were calculated to show the number of protein residues (Richardson et al., 2003). Finally, the electrostatic Poisson-Boltzmann (PB) potentials for all the structures were calculated using APBS molecular modelling software (DeLano Scientific LLC) implemented in PyMOL 0.99 (www.pymol.org) to define potential functional and interaction areas/clusters in the proteins. Potential values are given in units of kT per unit charge (k Boltzmann’s constant; T temperature).

## 3. Results and Discussion

### 3.1. ALDH gene families in the chickpea genome

We identified 27 unique ALDH gene sequences from the chickpea genome through database and bioinformatics searches. Information on the 27 chickpea sequences (name, locus ID, length, location on chromosome and features about the deduced peptide) is listed in Table 1. The exon number of the CaALDH genes ranged from 5 (NCBI: LOC101502106) to 21 (LOC101490622 and LOC101512568). The sizes of the deduced proteins varied markedly from 134 (LOC101502106) to 759 (LOC101512568) amino acids. The corresponding molecular weights varied from 15.07 to 82.38 kDa and the predicted isoelectric points (pIs) varied broadly from 4.34 to 9.49. As exhibited in other plant species, the wide range of pIs suggests that the chickpea ALDH proteins can work in various different subcellular environments, which is in accordance with the subcellular localization predicted for the sequences revealing that 44.4% (12 out 27) of CaALDHs can be localized to the cytoplasm (Supplementary Table 1). All 27 ALDH proteins contain a conserved ALDH domain (Pfam: PF00171) of variable length, which is a basic feature of ALDH families. The classification of protein families was made according to the criteria established by the ALDH Gene Nomenclature Committee (AGNC), namely the protein root symbol (ALDH) was followed by a family description number (1, 2, 3, etc.), a subfamily descriptor (A, B, C, etc.) and the individual gene number. As we used one gene model per locus, we did not include an extra lowercase letter to designate the number of variant. Thus, using the AGCN criteria, the ALDH proteins from chickpea fall into 10 families based on their sequence identities (Fig. 1). These families are also present in other vascular plants, suggesting that these 10 families may have evolved before the divergence of magnoliophyta and pteridophyta. Six chickpea families are represented by a single gene (ALDH5, ALDH6, ALDH7, ALDH11, ALDH12 and ALDH22), whereas the remaining four families contain multiple members (ALDH2, ALDH3, ALDH10 and ALDH18). Families ALDH5, 12 and 22 are also defined by a single gene in *Arabidopsis* as well as some other plant species. It has been proposed that these families represent constituted housekeeping ALDH genes, involved in preservation of nontoxic aldehyde levels and central plant metabolism (Jimenez-Lopez et al., 2016). The ALDH2 family, which is the largest ALDH family in plants contains 5 genes in chickpea. The ALDH3 family in chickpea is comparatively abundant containing the largest number of members (10 genes) described in plants to date. Thus, chickpea ALDH3 family may be functionally important carrying out additional stresses-responses proteins among ALDHs, enabling it to tolerate environmental stress such as salinity, drought through detoxification of molecules generated under these different stresses to maintain oxidative homeostasis. Four out of the 14 distinct ALDH families seem to be missed in the chickpea genome (ALDH19, ALDH21, ALDH23, and ALDH24), however they are represented in most of the plant species by a single gene. It has been proposed that families ALDH21, ALDH23 and ALDH24 played important roles in the transition of aquatic plants to terrestrial plants. Then, these families were lost during evolution of flowering plants (Rejeb et al., 2014; W. Wang et al., 2017). The family ALDH19 is unique among plants as only a single gene has been found in tomato, suggesting that this gene played an important role during evolution of that species (Jimenez-Lopez et al., 2016). This gene encodes a γ-glutamyl phosphate reductase, which catalyzes the reduction of l-glutamate 5-phosphate to 1-glutamate 5-semialdehyde (NADP-dependent) during the biosynthesis of proline from glutamate (García-Ríos et al., 1997).

**Table 1.** The aldehyde dehydrogenase gene superfamily in chickpea.

| Gene ID | Locus ID | Chromosome | Chr Start | Chr End | Strand | RNA accession | Exon number | Protein accession | Amino Acids number | Molecular Weight | Isoforms number | Isoelectric point |
| --- | --- | --- | --- | --- | --- | --- | --- | --- | --- | --- | --- | --- |
| CaALD H3F1 | LOC101 497113 | Ca1 | 1113 5019 | 1113 0555 | - | XM_004 486911 | 10 | XP_0044 86968 | 494 | 54.82 | 1 | 8.10 |
| CaALD H22A1 | LOC101 512347 | Ca1 | 1718 0053 | 1717 1549 | - | XM_004 487701 | 14 | XP_0044 87758 | 595 | 65.35 | 1 | 6.72 |
| CaALD H7A1 | LOC101 513733 | Ca1 | 2303 9421 | 2304 6624 | + | XM_012 718791 | 15 | XP_0125 74245 | 508 | 54.09 | 2 | 5.70 |
| CaALD H5F1 | LOC101 506901 | Ca1 | 3764 5851 | 3765 8363 | + | XM_004 488493 | 20 | XP_0044 88550 | 530 | 56.59 | 1 | 6.58 |
| CaALD H18B3 | LOC101 499756 | Ca3 | 8714 224 | 8700 487 | - | XM_012 713409 | 20 | XP_0125 68863 | 717 | 77.75 | 2 | 5.96 |
| CaALD H3H3 | LOC101 515558 | Ca4 | 3831 3842 | 3832 5387 | + | XM_004 498289 | 10 | XP_0044 98346 | 488 | 53.06 | 1 | 8.43 |
| CaALD H10A8 | LOC101 507930 | Ca5 | 3996 3842 | 3997 1506 | + | XM_004 501904 | 15 | XP_0045 01961 | 503 | 54.53 | 1 | 5.37 |
| CaALD H3H2 | LOC101 510937 | Ca5 | 4400 8221 | 4400 2223 | - | XM_004 502425 | 11 | XP_0045 02482 | 488 | 53.18 | 3 | 7.01 |
| CaALD H3H4 | LOC101 511680 | Ca5 | 4402 4284 | 4401 6817 | - | XM_004 502428 | 10 | XP_0045 02485 | 486 | 52.99 | 1 | 8.33 |
| CaALD H18B2 | LOC101 490622 | Ca6 | 1317 458 | 1311 762 | - | XM_012 716567 | 21 | XP_0125 72021 | 715 | 77.65 | 1 | 6.62 |
| CaALD H2C5 | LOC101 493969 | Ca6 | 3278 797 | 3283 156 | + | XM_004 503375 | 10 | XP_0045 03432 | 480 | 52.33 | 1 | 6.44 |
| CaALD H3H1 | LOC101 505038 | Ca6 | 6829 385 | 6835 166 | + | XM_004 503842 | 12 | XP_0045 03899 | 542 | 59.76 | 2 | 7.96 |
| CaALD H6B2 | LOC101 490310 | Ca6 | 1517 7302 | 1517 0648 | - | XM_004 504810 | 19 | XP_0045 04867 | 539 | 57.63 | 1 | 7.08 |
| CaALD H18B1 | LOC101 512568 | Ca6 | 4454 1903 | 4452 7947 | - | XM_027 335197 | 21 | XP_0271 90998 | 759 | 82.38 | 4 | 6.82 |
| CaALD H3F2 | LOC101 491914 | Ca6 | 5341 6538 | 5342 6529 | + | XM_004 507038 | 10 | XP_0045 07095 | 488 | 54.56 | 1 | 9.22 |
| CaALD H11A3 | LOC101 510843 | Ca7 | 1260 647 | 1264 733 | + | XM_004 507665 | 9 | XP_0045 07722 | 496 | 52.81 | 1 | 6.53 |
| CaALD H12A1 | LOC101 490107 | Ca7 | 8738 308 | 8744 740 | + | XM_004 508712 | 16 | XP_0045 08769 | 553 | 61.30 | 1 | 6.17 |
| CaALD H10A9 | LOC101 506136 | Ca7 | 9155 132 | 9150 438 | - | XM_004 508765 | 14 | XP_0045 08822 | 503 | 54.40 | 1 | 5.37 |
| CaALD H2B4 | LOC101 490532 | Ca7 | 9459 504 | 9464 830 | + | XM_004 508796 | 12 | XP_0045 08853 | 536 | 58.58 | 3 | 7.57 |
| CaALD H3F3 | LOC101 511819 | Ca7 | 1445 5263 | 1445 0584 | - | XM_012 718277 | 10 | XP_0125 73731 | 488 | 54.13 | 1 | 7.99 |
| CaALD H2B7 | LOC101 492709 | Ca7 | 2140 4965 | 2139 9791 | - | XM_004 509777 | 11 | XP_0045 09834 | 539 | 58.04 | 1 | 6.58 |
| CaALD<br>H2C6 | LOC101<br>513875 | Ca8 | 1499<br>2271 | 1498<br>3690 | - | XM_012<br>719313 | 9 | XP_0125<br>74767 | 498 | 44.10 | 1 | 5.55 |
| CaALD<br>H2C4 | LOC101<br>514219 | Ca8 | 1499<br>8332 | 1500<br>2867 | + | XM_004<br>512910 | 9 | XP_0045<br>12967 | 503 | 54.64 | 2 | 6.19 |
| CaALD<br>H3H7 | LOC101<br>502106 | Un | 0 | 0 | - | XM_004<br>514027 | 0 | XP_0045<br>14084 | 134 | 15.07 | 1 | 9.49 |
| CaALD<br>H18B4 | LOC105<br>852801 | Un | 0 | 0 | - | XM_012<br>719507 | 0 | XP_0125<br>74961 | 248 | 27.72 | 1 | 4.34 |
| CaALD<br>H3H5 | LOC101<br>497514 | Un | 0 | 0 | - | XM_027<br>330984 | 0 | XP_0271<br>86785 | 214 | 23.51 | 4 | 9.21 |
| CaALD<br>H3H6 | LOC101<br>488602 | Un | 0 | 0 | - | XM_027<br>330333 | 0 | XP_0271<br>86134 | 145 | 16.19 | 4 | 9.47 |

**Fig. 1.**
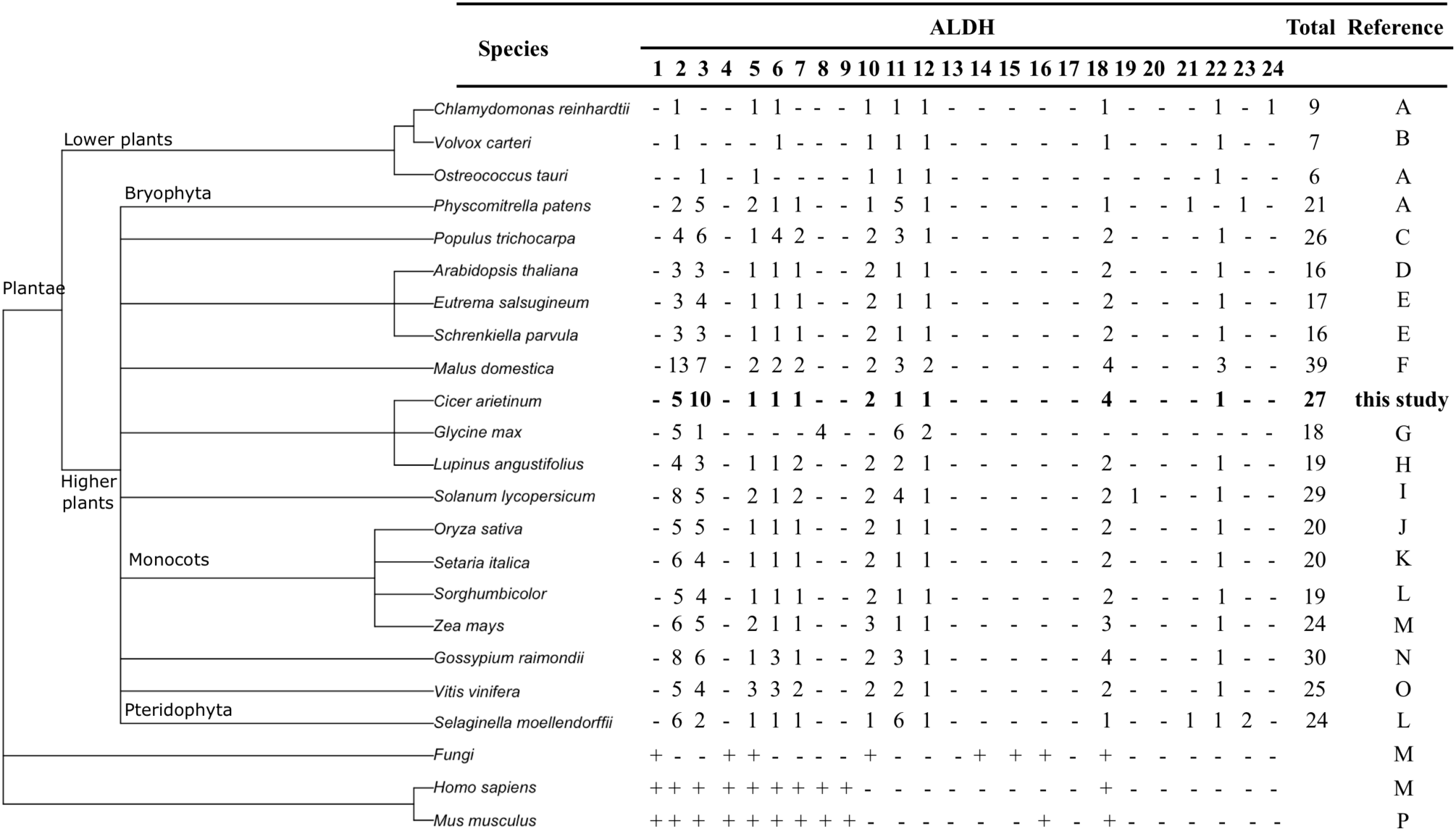
Distribution of ALDH families (1–24) in several species. The phylogenetic tree on the left, based on the taxonomic identifications of the species, was generated using the Taxonomy Common Tree Tools on the NCBI website (http://www.ncbi.nlm.nih.gov/guide/taxonomy/). The names of the ALDH families are listed on top of the table. The references are as follows: A, (Wood and Duff, 2009); B, (Prochnik et al., 2010); C, (Tian et al., 2015); D, (Kirch et al., 2004); E, (Hou and Bartels, 2015); F, (Li et al., 2013); G, (Kotchoni et al., 2012); H, (Jimenez-Lopez, 2016); I, (Jimenez-Lopez et al., 2016); J, (Gao and Han, 2009); K, (Chen et al., 2014); L, (Brocker et al., 2013); M, (He et al., 2014; Jimenez-Lopez et al., 2010); O, (He et al., 2014; Zhang et al., 2012); P, (L. Wang et al., 2017). Symbols + and − represent presence or absence, respectively.

Compared to other well-characterized plant ALDH families, such as *Arabidopsis*, maize, soybean, or rice, chickpea contains one of the most expanded ones, following the 39 ALDH genes in *M. domestica*, 30 in *Gossypium raimondii* and 29 in *Solanum lycopersicum*. Similar to *Gossypium* spp. (Guo et al., 2017; He et al., 2014), or *Oryza sativa* (Kotchoni et al., 2010), the four sequences of the chickpea ALDH18 family contain an AA-kinase domain, which is not found in other families, and lack the two other conserved sites (PS00687 and PS00070; Supplementary Table 1).

Finally, we mapped ALDH genes into chromosomes in order to gain an insight into the genome organization. Based on the available *C. arietinum* genome assembly, 23 out of the 27 CaALDH genes were distributed among seven of the eight chromosomes. We could not map LOC101502106, LOC101497514, LOC101488602 (members of ALDH3 subfamily), and LOC105852801 (ALDH18B4). The other 23 ALDH genes were unevenly distributed through the chickpea genome. Two chromosomes contained the highest number with six ALDH genes (chromosome 6 and 7), whereas chromosome 3 and 4 contained one ALDH gene, respectively. Chromosome 8, which is the shortest in the chickpea genome, contained two ALDH genes (LOC101513875 and LOC101514219). No ALDH gene could be found in the chromosome 2 (Fig. 2).

**Fig. 2.**
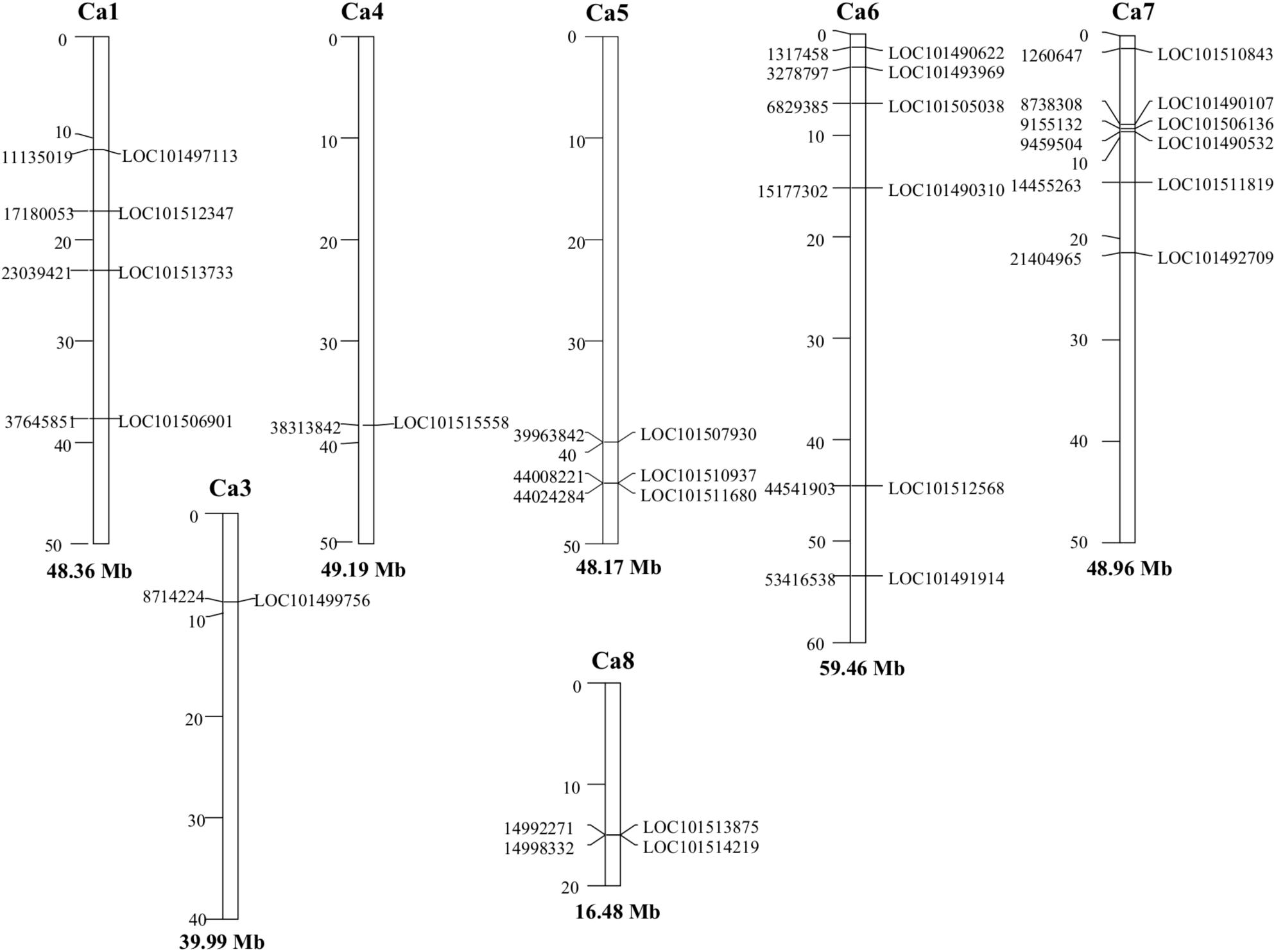
Genomic distribution of ALDH genes on chickpea chromosomes. Only those chromosomes bearing CaALDH genes are represented. The chromosome numbers and sizes (Mb) are indicated at the top and bottom of each bar, respectively.

### 3.2. Evolutionary relationships of ALDH gene families between chickpea and *Medicago truncatula*

In order to explore the evolution of the CaALDH genes, we compared the syntenic blocks of the chickpea and the model legume *Medicago truncatula* genomes. In previous studies, synteny analyses have revealed extensive conservation and good collinearity between both legumes (Gupta et al., 2017; Parween et al., 2015). In the current study, we identified large-scale synteny blocks containing orthologues from six ALDH families (ALDH6, ALDH7, ALDH11, ALDH12, ALDH18, and ALDH22), including 8 CaALDH genes from chickpea and 8 ALDH genes from *Medicago* (Supplementary Table 2). Five pairs of orthologous genes appeared to be single chickpea-to-*Medicago* ALDH gene correspondences. It is likely that these genes/families derived from a common ancestor of chickpea and *Medicago* conserved during evolution. Furthermore, we also found instances of a single chickpea gene corresponded to multiple *Medicago* genes, in addition to several chickpea duplications corresponded to a single *Medicago* gene. The remaining four chickpea families (ALDH2, ALDH3, ALDH5, and ALDH10) could not be mapped to any syntenic block.

### 3.3. Phylogenetic analysis of chickpea ALDH genes

To study the evolutionary relationship of the ALDH gene superfamily among different species, a phylogenetic tree was generated with full-length of 79 well-characterized ALDH proteins from *Arabidopsis, Glycine max* and *Zea mays* (Fig. 3). The result was consistent with previous findings (Chen et al., 2014; Guo et al., 2017; Jiang et al., 2019), and showed that different family proteins in the same species did not clustered together. However, it grouped the same family proteins of different species, and every cluster containing both, monocotyledon and dicotyledon ALDH gene members. The ALDH19 family is not included here as our analyses did not incorporate any sequence from tomato (Li et al., 2013). The phylogenetic tree indicates that most of the ALDH families represent a common plant ALDH core (ALDH2, ALDH3, ALDH5, ALDH6, ALDH10, ALDH11, ALDH22). The ALDH18 family is the most phylogenetically distant group related to the remaining families, indicating that these proteins had the greatest degree of sequence divergence from the other ALDH families and did not contain the conserved ALDH active sites (Brocker et al., 2013). The majority of CaALDHs grouped more closely related to soybean than to *Arabidopsis* or maize, which is consistent with the evolutionary relationships among the four species. It is worth mentioning that families ALDH5, ALDH6 and ALDH12, all of them represented by only one chickpea sequence, grouped with ALDH proteins from *Arabidopsis* and maize, but the clusters did not contain any soybean sequence (Fig. 1). These sequences likely were independently lost in the soybean genome. On the contrary, the cluster grouping family ALDH7, represented by only one chickpea member, contained four soybean sequences. In this case, the soybean genome most likely increased this ALDH family by duplication events.

**Fig. 3.**
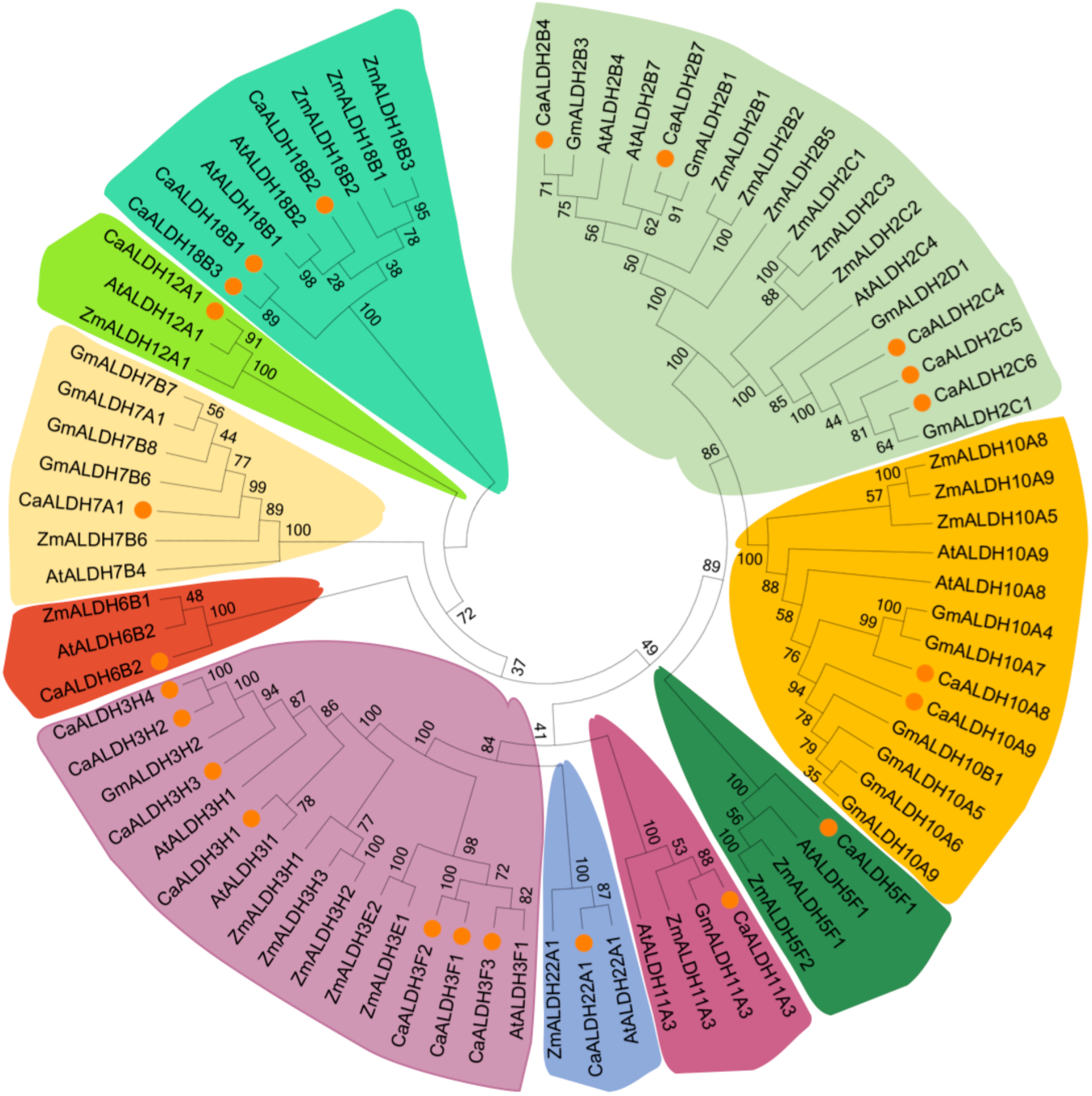
Phylogenetic tree of ALDH proteins from chickpea, *G. max* (Gm), *A. thaliana* (At), and *Zea mays* (Zm). Alignment of 79 ALDH protein sequences from four plant species was conducted with ClustalW2, and the phylogenetic tree was constructed using MEGA 6 based on the Neighbor-joining (NJ) method. Bootstrap values in percentage (500 replicates) are labelled on the nodes. CaALDHs are marked with solid orange circles.

It is clear that cluster with family ALDH3 was mostly made because of the remarkable expansion of this family in the chickpea genome. In *Arabidopsis*, the expression of class3 ALDHs is induced by environmental stresses such as drought, salinity, ABA exposure, heavy metals and pesticides (Kirch et al., 2005; Kotchoni et al., 2006; Missihoun et al., 2011; Stiti et al., 2011). The notable expansion of the CaALDH3 gene families compared with other plant species suggests that these ALDH genes may be essential for chickpea to cope with environmental stresses.

### 3.4. ALDH expansion: gene duplications

The expansion of gene families is based on gene duplications, which in turn, mainly rely on segmental and tandem duplications (Cannon et al., 2004). Based on a comprehensive analysis of chromosomal locations and sequence similarities, 59.3% in 16 out of 27 ALDH sequences, ALDH genes appear to be associated with either local duplication events or duplications to unlinked *loci* (Fig. 4). There is no support for tandemly duplicated ALDH genes in the genome of Chinese cabbage (Jiang et al., 2019), however tandem duplications have been shown to occur in the ALDH family of grapes, apples, and soybeans (Li et al., 2013; W. Wang et al., 2017; Zhang et al., 2012), as well as the monocot species rice and millet (Chen et al., 2014; Gao and Han, 2009). Chickpea ALDH genes mapped on the same chromosomes are candidates to have undergone local gene duplications. We found two genes on chromosome 5 (LOC101510937, LOC101511680) and two genes on chromosome 8 (LOC101513875, LOC101514219) that met the criteria to form a cluster as described in subsection 2.2. These two pair of genes are separated by <10 kb, respectively. The other duplicated genes (75% duplications) are located on different chromosomes, suggesting that segmental duplications played a major role in the expansion of the ALDH gene family in chickpea.

**Fig. 4.**
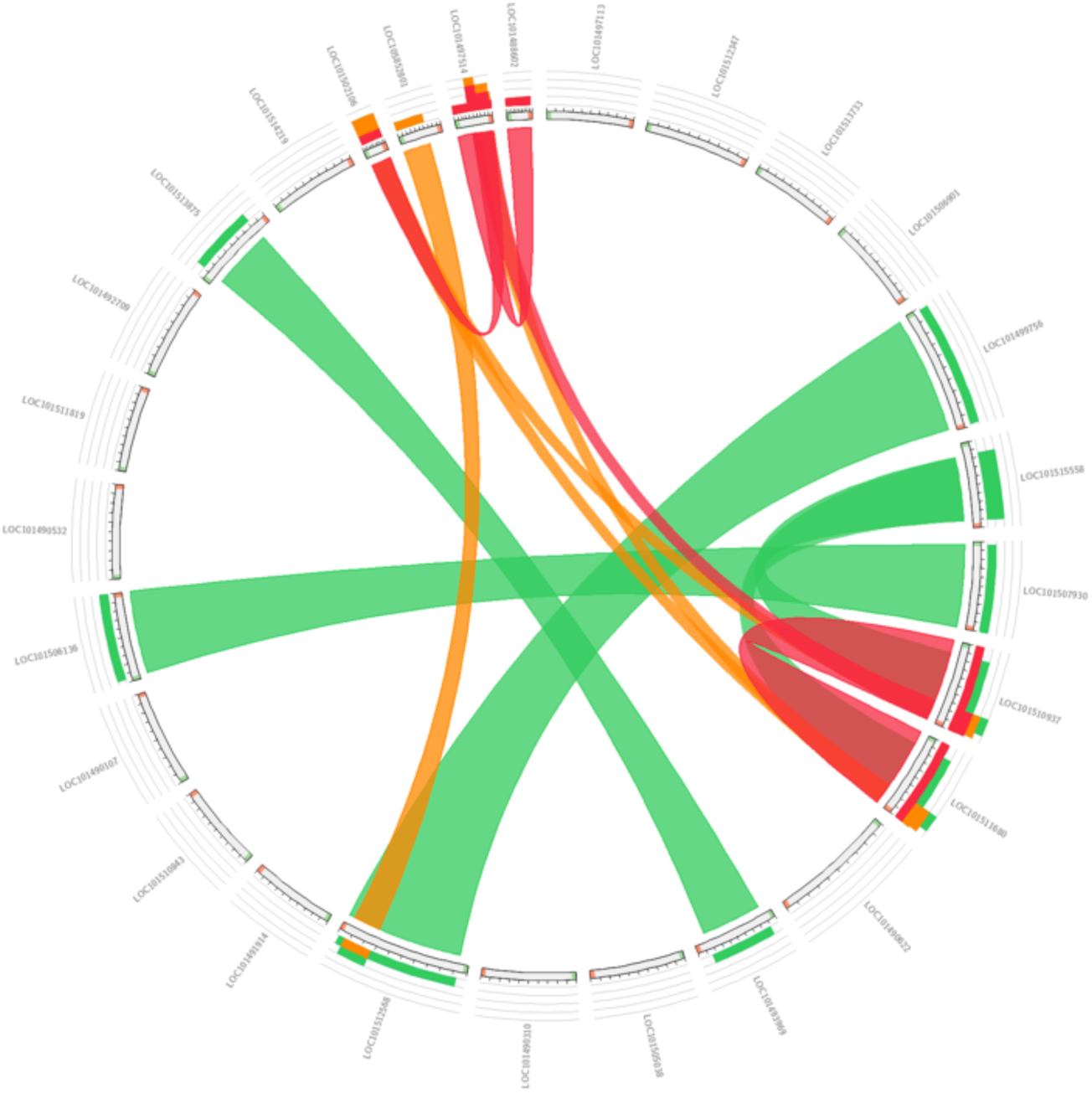
Gene duplications. Similarity of ALDH genes. Red colour shows the highest similarity (> 95% identity), followed by orange (90-95%) and green (80-90%) colours.

### 3.5. Expression profiles of CaALDH genes

In order to gain an insight into the functional roles of the ALDH genes, we analyzed their expression patterns in different tissues using available EST datasets (EST, 2017). Considering the stringent criterion described in subsection 2.4., 11 ALDH genes had expression support (26 ESTs). One ALDH gene (LOC101506136) hit 8 ESTs, whereas LOC101510843 and LOC101490310 hit four and three ESTs, respectively (Supplementary Table 3). Regarding the plant tissues, root tissue was the most common hit (18 hits) followed by leaves (5 hits). The experimental conditions of these libraries suggest a role in a variety of environmental responses by the ALDH superfamily. Most of the libraries were constructed in response to drought stress (14 hits) but we also found ESTs from libraries in responses to insect attack, Cd toxicity, and response to thidiazuron, a synthetic plant regulator of morphogenetic processes that induces the expression of stress-related genes (Dewir et al., 2018; Zhang et al., 2006). Interestingly, 5 ESTs were found in specific subtracted cDNA libraries from infected roots with the soil-borne fungus *Fusarium oxysporum*, which is a serious threat to chickpea production. Based on this finding, we aimed to gain an insight into the role of the ALDHs in the *Fusarium* wilt resistance by using Magic-BLAST, a novel tool allowing the mapping of large next-generation sequencing runs against a reference database (Boratyn et al., 2019). From publicly available transcriptome datasets, we selected two libraries constructed with the chickpea genotype WR315, as this genotype is commonly used as a parental line in the breeding program. Some sequences showed extremely low ALDH count numbers, suggesting that they are expressed at very low levels in root tissues (LOC101497113, LOC101491914, and LOC101511819). Overall, the ALDH counts are highly correlated between non-inoculated and inoculated plants (R = 0.91). However, two genes were more abundant in a given condition: LOC101510843 (*CaALDH11A3*) was over-represented in roots of control plants, while LOC101510937 (*CaALDH3H2*) showed a larger count numbers in inoculated plants (Fig. 5).

**Fig. 5.**
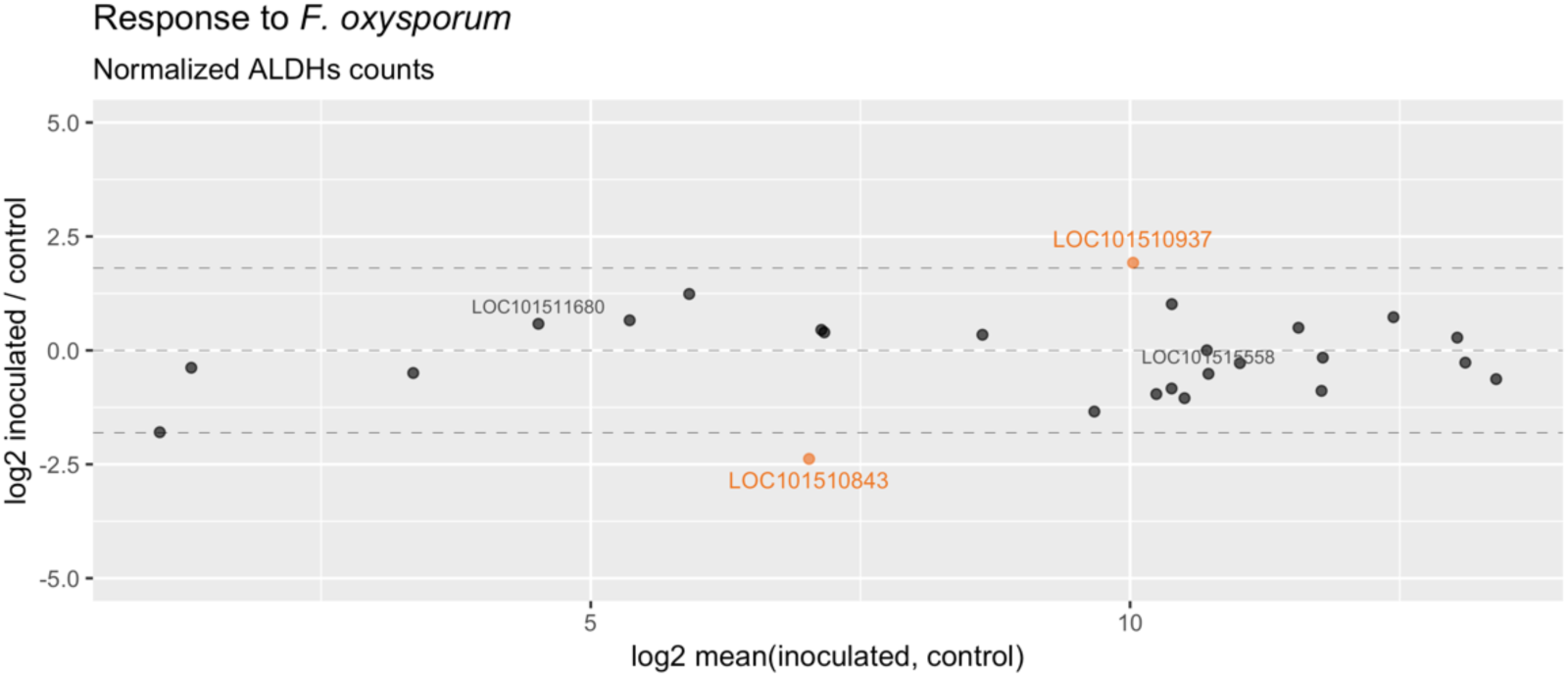
Gene expression analysis in response to *Fusarium oxysporum*.MA-plot of mean expression signal *vs* log2-normalized counts of ALDH genes in two chickpea transcriptome libraries (inoculated *vs* control). Genes highly enriched (counts ratio >3.5-fold) in any of the conditions are shown in orange colour, whereas the duplicated genes LOC101511680 and LOC101515558 are shown in grey colour.

The ALDH11 family has a crucial function in the generation of NADPH, which is the main source for mannitol biosynthesis in many plant species (Plaxton, 1996). The duplication of ALDH11 genes in tomato has been related to an increase in sugar production (Jimenez-Lopez et al., 2016). On the other hand, the phylogenetic tree showed that the chickpea ALDH3H2 is closely related to the *Arabidopsis* ALDH3H1 (AT1G44170, NP_175081), which is a protein localized in the cytoplasm involved in responses to desiccation, response to abscisic acid stimulus, and induced at high salt concentrations (Hou and Bartels, 2015). Interestingly, LOC101515558 and LOC101511680 (*ALDH3H3* and *ALDH3H4*, respectively), which are duplicated with LOC101510937 (Fig. 4) show a different expression pattern, as they are not differentially abundant in any condition. This result suggests that CaALDH3H2 and the duplicated sequences CaALDH3H3, and CaALDH3H4 are probably regulated in different ways. The remarkable expansion of the ALDH3 family in chickpea may have evolved as a consequence of functional specialization. LOC101510937 is a particularly valid candidate for further experimental validation.

### 3.6. Structure-based functional analysis of selected members of the ALDH3 family

Crystallographic structures of particular ALDH proteins have greatly facilitate the correlative study between structure and functionality of many ALDH in different organisms (Jimenez-Lopez et al., 2010; Kotchoni et al., 2010). To the best of our knowledge, structure-functional characterization and 3D comparative analysis of different members of the ALDH protein superfamily have been only performed in few organisms such as rice (Kotchoni et al., 2010), maize (Jimenez-Lopez et al., 2010), tomato (Jimenez-Lopez et al., 2016) or lupin (Jimenez-Lopez, 2016).

Through computational homology modelling, we could uncover the characteristics 2D and 3D structure features for the catalytic active sites and the NAD(P)^+^ - binding clefts of representative members of the ALDH3 family (Fig. 6). We addressed an accurate structural assessment of the models by comparative analysis with their best templates (PDBs accession numbers 1ad3, 5nno, 5ucd, and 4qgk, respectively). The stereo-chemical and energy minimization parameters displaying the following data: the analysis of the best templates showed z-value (normalized QMEAN4 scores) of -0.1, 0.23, 1.16 and 0.66 for the Q-mean parameter (estimating the model reliability ranging between 0 and 1, respectively, and 0.94, 0.35, and 0.22 for ALDH3 models sequences XP_004502485, XP_004502482, and XP_004498346, respectively. These z-scores are within good values expected for these models in comparison to the best templates used for building the structures. An additional parameter to check the overall quality of the structures, ProSA, showed a z-score of −9.53, −8.16, and −8.94 for ALDH3 models, and, −10.31, -9.72, -10.67 and -9.58 for the individual crystallographic structural templates. Both, Q-mean and ProSA parameters show similar values for the ALDH3 models in comparison to the PDB structures. This indicates that the ALDH3 protein models are accurate and close to their templates in structure quality. Thus, the analysis of the stereochemistry of the model using Procheck analysis shows that 93.7 %, 98%, 97.7, and 98.1% of the structural residues locate in favourable regions in the respective templates; 6, 1.8, 2.1, and 1.6 in allowed regions, 0.3, 0.2, 0.2 and 0.2 in generally allowed regions; 0, 0, 0 and 0% for the four templates in disallowed regions. The same values for the ALDH3 models were 94, 5, 0.6, and 0.4%; 93.6, 4.7, 1.6, and 0.1%; and 96.7, 2.5, 0.8, 0% from the sequences XP_004502485, XP_004502482, and XP_004498346, respectively, finding even more residues located in favourable regions, less residues in allowed regions, and a similar situation in generally allowed and non-favourable regions.

**Fig. 6.**
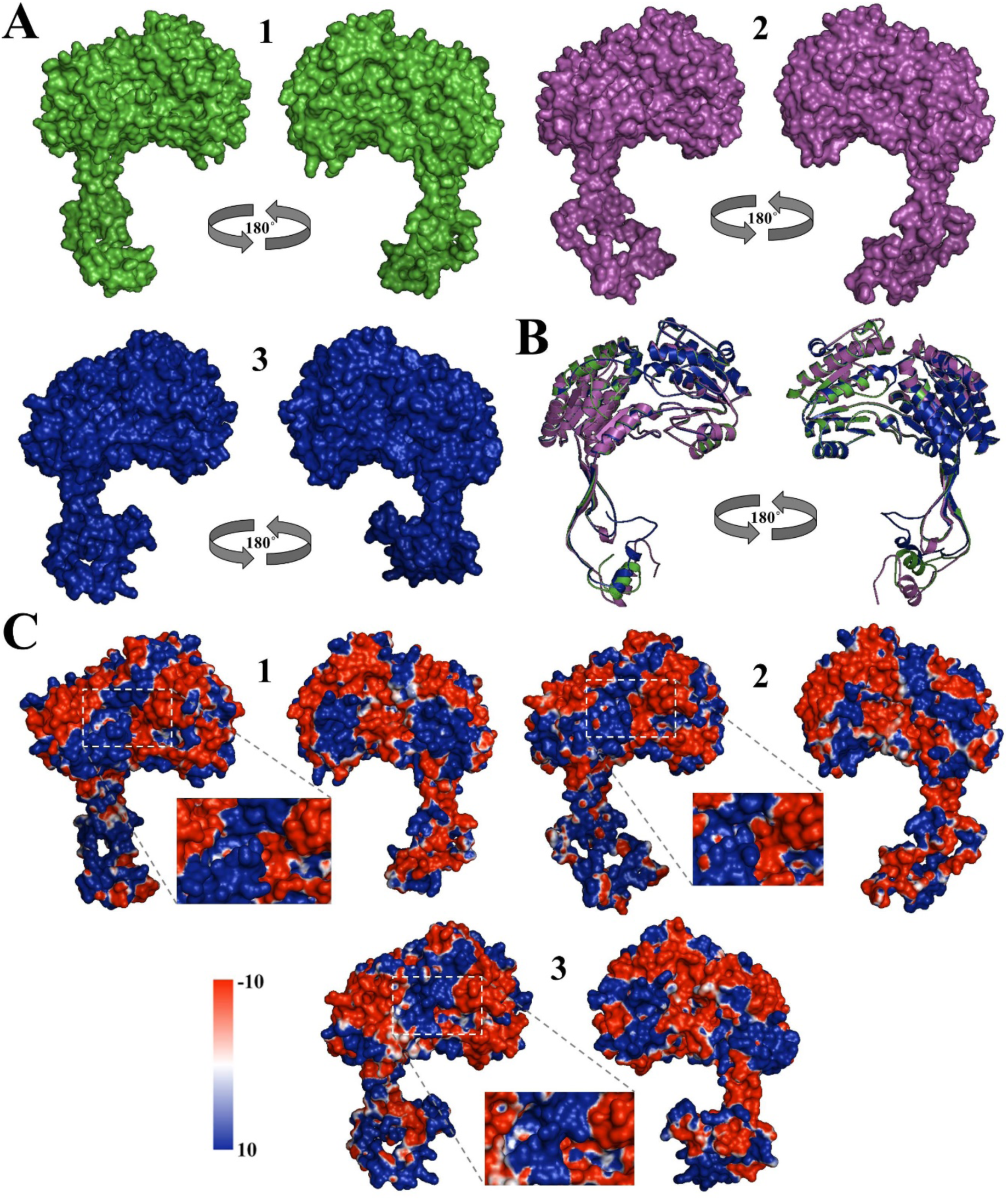
Structural analysis of *C. arietinum* ALDH3 superfamily. (A) Three-dimensional structure of chickpea ALDH3 members (1) XP_004502485, (2) XP_004502482 (2), and (3) XP_004498346). Structures were depicted as surface diagram. Two views rotated 180° around the X-axis are provided. (B) Superimposition between chickpea ALDH3 members (1) XP_004502485, (2) XP_004502482, and (3) XP_004498346 depicted in green, purple and blue colours, respectively. (C) Electrostatic potential representation on the ALDH3 protein surfaces, showing a detailed view of the coenzyme-catalytic domain. The surface colours are clamped at red (-10) or blue (+10). Two views rotated 180° around the X-axis are provided for members of the ALDH3 superfamily.

Based on all the above parameters, we confirmed the accuracy and reliability of the structural models built for the ALDH3 sequences on the basis of their crystallographic templates. Therefore, the protein models could be used for further structure-functional analyses, which revealed important differences. Specifically, the structure of the oligomerization region, where 2D (α-helices and β-sheets) number, curvature angle, and length, as well as folding characteristics were prominent. Although the overall topology of the ALDH surface structures were comparable among the sequences, XP_004498346 exhibited loop bigger than the other structures, whereas its coenzyme binding region seems to be narrower and less open to the surface (Fig. 6A). Since the oligomerization domain seems to be a main feature differentiating the three protein models, this may highlight differential functionality since some ALDH are only active as a dimer (Keller et al., 2014, 2010). The comparison of the represented cartoon 2D elements between the three models by superimposition of the three ALDH3 structures is shown in Fig. 6B. We found some differences between the coils structures, particularly in the oligomerization domain, and the N-terminal region, which is longer in XP_004498346. This particular feature may confer protection from the solvent surface, thus smaller macromolecules would be more accessible to the co-enzyme and catalytic domains.

The analysis of the surface distribution of charges showed a comparable pattern between XP_004502485, and XP_004502482, whereas the electrostatic potential in XP_004498346 was dominated by positive charges along catalytic and oligomerization domains (Fig. 6C). We can hypothesize that the pattern of charges distribution may indicate differences in the possible functional mechanism and/or interaction with other proteins and subcellular localization leading to differential functional activities.

Particular characteristics of binding clefts exhibit certain degree of conformational flexibility for cofactors highlighting functional dynamic preference for the oxidized or reduced NADH/NAD^+^ (Jimenez-Lopez, 2016; Jimenez-Lopez et al., 2016, 2010). In our study, the analysis shows residues included in the catalytic cleft (substrate and cofactor binding sites) with conserved basic amino acid N residue in different positions and located in the opposite side of the NAD^+^ ring, and other conserved C residues (Fig. 7). These residues may be involved in the enzymatic mechanism of the ALDH, with the characteristic nucleophilic attack and proton abstraction from the C residue during the course of the reaction, and with influence in the thiol extraction step during the catalysis reaction by ALDHs (Perez-Miller and Hurley, 2003). The alignment of the three ALDH sequences display the residues implicated in the cofactor and catalytic binding cleft, and the functional catalytic residues directly involved in the catalytic reactions (Fig. 7). Functional analysis of the three ALDH3 members shows a co-enzymatic and catalytic environment well conserved with residues that maintain these two functional clefts, including a W126 and H303 binding to NAD^+^ phosphate, E153 interacting to pentose ring and F353 to nicotinic ring; a coenzyme - catalytic binding domain integrated by L189, T186, I207, Y199, K150, L132, A125, S152, W126, N127, E153, G200, N201, G181, V183, V148, N302, R306, H303, E350, E351, F353, and important amino acids involved in the catalytic reaction such as C257 and N201 (Fig. 7A and 7C). Based in the distance from the cofactor to the C257, the residue E may be the functional one, where the putative mechanism for this ALDH would consist in the activation of C257 by a base (possibly N201), starting a nucleophilic attack on the carbonyl carbon of the aldehyde (Fig. 7B). N201 is in charge of the correct positioning of the polar aldehyde head group in order to liberate a hydride ion, which in turn is transferred to NAD. Finally, a water molecule transfers a proton to initiate a nucleophilic attack on the carbonyl carbon of the covalently bound substrate. Thus, an oxyanion breaks the thio-hemiacetal bond and releases the product (Lloyd et al., 2007).

**Fig. 7.**
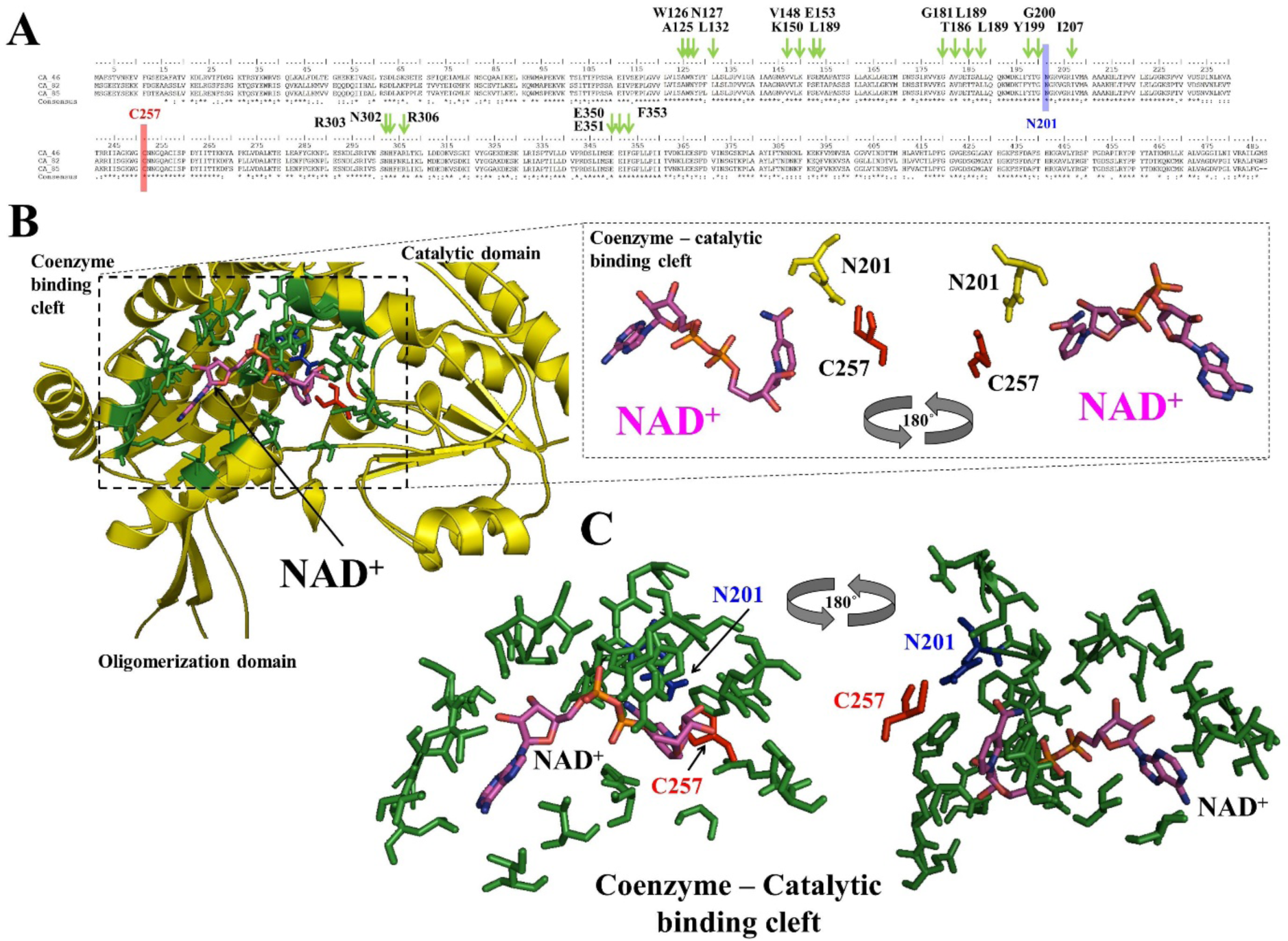
Coenzyme and ligand-binding domain analysis of ALDH3 superfamily. **(A)** Sequence alignment of the ALDH3 members C_85 (XP_004502485), C_82 (XP_004502482), and C-46 (XP_004498346). Catalytic residues are highlighted in red (C257) and blue (N201) colours. Residues involved in the coenzyme binding cleft are highlighted in green colour. All these residues are 100% conserved between these three members of the ALDH3 superfamily. (B) Distribution of the NADH cofactor and the spatial distribution of the residues that integrate the cofactor-substrate binding cleft (green colour) and integrated in the carton representation of the XP_004502485 ALDH3 structure. Residues are depicted as stick and green coloured and red/blue colours the catalytic residues. A Detailed representation of the amino acids interacting with NADH corresponding to the catalytic residues C257 and N201. (C) Detailed view of the coenzyme-catalytic binding cleft and the spatial distribution of the residues involved in building the cleft. Residues are depicted as stick and coloured in green colour. Critical catalytic amino acids (C257 and N201) are highlighted in red and blue background, respectively. Two views rotated 180° around the x-axis are provided for the coenzyme-catalytic binding cleft of the ALDH3 member XP_004502485.

## 4. Conclusions

1. Plants are continuously exposed to different types of abiotic and biotic stresses. Plant molecular responses induce the generation of reactive oxygen species, which will in turn interfere with cell structure and metabolic balance in cells. To protect themselves, plants produce stress-responsive proteins, such as ALDHs that contribute to aldehyde homeostasis as scavengers to eliminate toxic aldehydes.
2. In the present study, performing a series of comprehensive analyses including chickpea genome analysis, ALDH genes identification and naming, comparative phylogeny, ALDH genes expression profiles assessment and structure-based functional analysis, we identified 27 unique ALDH sequences in the chickpea genome.
3. Most of the sequences were originated from duplication events. Chickpea exhibits a remarkable expansion in the ALDH3 family, showing the largest number of members compared to any other plant species described so far.
4. The expression results give consistent support in the functional roles of the ALDH genes, mostly being involved in responses to desiccation and drought conditions, but also responses to biotic stress. Furthermore, the expression data revealed that some of the duplicated members in a group exhibited different expression patterns, suggesting that functional diversification is a feature in the evolution of these genes.
5. The modelling of some ALDH3 family members on the basis of crystallographic protein structures provided insights into the relationship between interactive surfaces and the ALDH catalytic domains; setting structural features of binding pockets and the specificity of cofactor NAD and substrate within a particular family of ALDH.
6. Based on expression data support and close phylogenetic relationships with other well-characterized proteins, some chickpea ALDHs (such as LOC101510937) are good candidates for further characterization. These candidates may become targets for improving chickpea adaptation to adverse environmental or biotic stresses in the breeding program. Furthermore, our study also provides a foundation for further comparative genomic analyses and a framework to trace the dynamic evolution of the ALDH superfamily.

## Author Contribution

Conceptualization, JVD and JG; Formal analysis, RCM, T.M, JCJL, and JVD; Funding acquisition, TM; Writing—original draft, RCM, and JVD.; Writing—review & editing, JCJL, TM, JG, and JVD; All authors read and approved the manuscript.

## Data Availability Statement

All data generated or analyzed during this study are included in this article and its supplementary information files.

## References

Almagro Armenteros, J.J., Sønderby, C.K., Sønderby, S.K., Nielsen, H., Winther, O., 2017. DeepLoc: prediction of protein subcellular localization using deep learning. Bioinformatics 33, 3387–3395. doi: 10.1093/bioinformatics/btx431

Altschul, S.F., Gish, W., Miller, W., Myers, E.W., Lipman, D.J., 1990. Basic local alignment search tool. J. Mol. Biol. 215, 403–410. doi: 10.1016/S0022-2836(05)80360-2

Ameline-Torregrosa, C., Wang, B.-B., O’Bleness, M.S., Deshpande, S., Zhu, H., Roe, B., Young, N.D., Cannon, S.B., 2008. Identification and characterization of nucleotide-binding site-leucine-rich repeat genes in the model plant *Medicago truncatula*. Plant Physiol. 146, 5–21. doi: 10.1104/pp.107.104588

Bartels, D., 2001. Targeting detoxification pathways: an efficient approach to obtain plants with multiple stress tolerance? Trends Plant Sci. 6, 284–286.

Black, W., Vasiliou, V., 2009. The aldehyde dehydrogenase gene superfamily resource center. Hum Genomics 4, 136–142. doi: 10.1186/1479-7364-4-2-136

Boratyn, G.M., Thierry-Mieg, J., Thierry-Mieg, D., Busby, B., Madden, T.L., 2019. Magic-BLAST, an accurate RNA-seq aligner for long and short reads. BMC Bioinformatics 20, 405. doi: 10.1186/s12859-019-2996-x

Brocker, C., Vasiliou, M., Carpenter, S., Carpenter, C., Zhang, Y., Wang, X., Kotchoni, S.O., Wood, A.J., Kirch, H.-H., Kopečný, D., Nebert, D.W., Vasiliou, V., 2013. Aldehyde dehydrogenase (ALDH) superfamily in plants: gene nomenclature and comparative genomics. Planta 237, 189–210. doi: 10.1007/s00425-012-1749-0

Cannon, S.B., Mitra, A., Baumgarten, A., Young, N.D., May, G., 2004. The roles of segmental and tandem gene duplication in the evolution of large gene families in *Arabidopsis thaliana*. BMC Plant Biol. 4, 10. doi: 10.1186/1471-2229-4-10

Chen, Z., Chen, M., Xu, Z., Li, L., Chen, X., Ma, Y., 2014. Characteristics and expression patterns of the aldehyde dehydrogenase (ALDH) gene superfamily of foxtail millet (*Setaria italica* L.). PLoS ONE 9, e101136. doi: 10.1371/journal.pone.0101136

Claros, M.G., Vincens, P., 1996. Computational method to predict mitochondrially imported proteins and their targeting sequences. Eur. J. Biochem. 241, 779–786. doi: 10.1111/j.1432-1033.1996.00779.x

Darzentas, N., 2010. Circoletto: visualizing sequence similarity with Circos. Bioinformatics 26, 2620–2621. doi: 10.1093/bioinformatics/btq484

Dewir, Y.H., Nurmansyah Naidoo, Y., Teixeira da Silva, J.A., 2018. Thidiazuron-induced abnormalities in plant tissue cultures. Plant Cell Rep. 37, 1451–1470. doi: 10.1007/s00299-018-2326-1

Die, J.V., 2018. refseqR: Common computational operations working with GenBank. Zenodo. doi: 10.5281/zenodo.1188462

Emanuelsson, O., Nielsen, H., von Heijne, G., 1999. ChloroP, a neural network-based method for predicting chloroplast transit peptides and their cleavage sites. Protein Sci. 8, 978–984. doi: 10.1110/ps.8.5.978

EST, National Library of Medicine (US), National Center for Biotechnology Information, 2018. Available online: https://www.ncbi.nlm.nih.gov/est/ (accessed on Apr 10, 2019).

FAOSTAT, 2018. FAOSTAT. Crop Statistics. Available online http://faostat.fao.org (accessed Nov 20.18)

Gao, C., Han, B., 2009. Evolutionary and expression study of the aldehyde dehydrogenase (ALDH) gene superfamily in rice (*Oryza sativa*). Gene 431, 86–94. doi: 10.1016/j.gene.2008.11.010

García-Ríos, M., Fujita, T., LaRosa, P.C., Locy, R.D., Clithero, J.M., Bressan, R.A., Csonka, L.N., 1997. Cloning of a polycistronic cDNA from tomato encoding gamma-glutamyl kinase and gamma-glutamyl phosphate reductase. Proc Natl Acad Sci USA 94, 8249–8254. doi: 10.1073/pnas.94.15.8249

Guo, X., Wang, Yuanyuan, Lu, H., Cai, X., Wang, X., Zhou, Z., Wang, C., Wang, Yuhong, Zhang, Z., Wang, K., Liu, F., 2017. Genome-wide characterization and expression analysis of the aldehyde dehydrogenase (ALDH) gene superfamily under abiotic stresses in cotton. Gene 628, 230–245. doi: 10.1016/j.gene.2017.07.034

Gupta, S., Nawaz, K., Parween, S., Roy, R., Sahu, K., Kumar Pole, A., Khandal, H., Srivastava, R., Kumar Parida, S., Chattopadhyay, D., 2017. Draft genome sequence of *Cicer reticulatum* L., the wild progenitor of chickpea provides a resource for agronomic trait improvement. DNA Res. 24, 1–10. doi: 10.1093/dnares/dsw042

He, D., Lei, Z., Xing, H., Tang, B., 2014. Genome-wide identification and analysis of the aldehyde dehydrogenase (ALDH) gene superfamily of *Gossypium raimondii*. Gene 549, 123–133. doi: 10.1016/j.gene.2014.07.054

Hobohm, U., Sander, C., 1995. A sequence property approach to searching protein databases. J. Mol. Biol. 251, 390–399. doi: 10.1006/jmbi.1995.0442

Hou, Q., Bartels, D., 2015. Comparative study of the aldehyde dehydrogenase (ALDH) gene superfamily in the glycophyte *Arabidopsis thaliana* and *Eutrema halophytes*. Ann. Bot. 115, 465–479. doi: 10.1093/aob/mcu152

Jain, M., Misra, G., Patel, R.K., Priya, P., Jhanwar, S., Khan, A.W., Shah, N., Singh, V.K., Garg, R., Jeena, G., Yadav, M., Kant, C., Sharma, P., Yadav, G., Bhatia, S., Tyagi, A.K., Chattopadhyay, D., 2013. A draft genome sequence of the pulse crop chickpea (*Cicer arietinum* L.). Plant J. 74, 715–729. doi: 10.1111/tpj.12173

Jakoby, W.B., Ziegler, D.M., 1990. The enzymes of detoxication. J. Biol. Chem. 265, 20715–20718.

Jiang, X., Ren, J., Ye, X., Liu, M., Li, Q., Wang, L., Liu, Z., 2019. Genome-wide identification and analysis of the aldehyde dehydrogenase gene superfamily in Chinese cabbage (*Brassica rapa* L. ssp. *pekinensis*). Can. J. Plant Sci. doi: 10.1139/CJPS-2018-0205

Jimenez-Lopez, J.C., 2016. Narrow-leafed lupin (*Lupinus angustifolius* L.) functional identification and characterization of the aldehyde dehydrogenase (ALDH) gene superfamily. Plant Gene 6, 67–76. doi: 10.1016/j.plgene.2016.03.007

Jimenez-Lopez, J.C., Gachomo, E.W., Seufferheld, M.J., Kotchoni, S.O., 2010. The maize ALDH protein superfamily: linking structural features to functional specificities. BMC Struct. Biol. 10, 43. doi: 10.1186/1472-6807-10-43

Jimenez-Lopez, J.C., Lopez-Valverde, F.J., Robles-Bolivar, P., Lima-Cabello, E., Gachomo, E.W., Kotchoni, S.O., 2016. Genome-Wide Identification and Functional Classification of Tomato (*Solanum lycopersicum*) Aldehyde Dehydrogenase (ALDH) Gene Superfamily. PLoS ONE 11, e0164798. doi: 10.1371/journal.pone.0164798

Keller, M.A., Watschinger, K., Golderer, G., Maglione, M., Sarg, B., Lindner, H.H., Werner-Felmayer, G., Terrinoni, A., Wanders, R.J.A., Werner, E.R., 2010. Monitoring of fatty aldehyde dehydrogenase by formation of pyrenedecanoic acid from pyrenedecanal. J. Lipid Res. 51, 1554–1559. doi: 10.1194/jlr.D002220

Keller, M.A., Zander, U., Fuchs, J.E., Kreutz, C., Watschinger, K., Mueller, T., Golderer, G., Liedl, K.R., Ralser, M., Kräutler, B., Werner, E.R., Marquez, J.A., 2014. A gatekeeper helix determines the substrate specificity of Sjögren-Larsson Syndrome enzyme fatty aldehyde dehydrogenase. Nat. Commun. 5, 4439. doi: 10.1038/ncomms5439

Kirch, H.-H., Bartels, D., Wei, Y., Schnable, P.S., Wood, A.J., 2004. The ALDH gene superfamily of *Arabidopsis*. Trends Plant Sci. 9, 371–377. doi: 10.1016/j.tplants.2004.06.004

Kirch, H.-H., Schlingensiepen, S., Kotchoni, S., Sunkar, R., Bartels, D., 2005. Detailed expression analysis of selected genes of the aldehyde dehydrogenase (ALDH) gene superfamily in *Arabidopsis thaliana*. Plant Mol. Biol. 57, 315–332. doi: 10.1007/s11103-004-7796-6

Kotchoni, S.O., Jimenez-Lopez, J.C., Gao, D., Edwards, V., Gachomo, E.W., Margam, V.M., Seufferheld, M.J., 2010. Modeling-dependent protein characterization of the rice aldehyde dehydrogenase (ALDH) superfamily reveals distinct functional and structural features. PLoS ONE 5, e11516. doi: 10.1371/journal.pone.0011516

Kotchoni, S.O., Jimenez-Lopez, J.C., Kayodé, A.P.P., Gachomo, E.W., Baba-Moussa, L., 2012. The soybean aldehyde dehydrogenase (ALDH) protein superfamily. Gene 495, 128–133. doi: 10.1016/j.gene. 2011.12.035

Kotchoni, S.O., Kuhns, C., Ditzer, A., Kirch, H.-H., Bartels, D., 2006. Over-expression of different aldehyde dehydrogenase genes in *Arabidopsis thaliana* confers tolerance to abiotic stress and protects plants against lipid peroxidation and oxidative stress. Plant Cell Environ. 29, 1033–1048. doi: 10.1111/j.1365-3040.2005.01458.x

Lee, T.-H., Tang, H., Wang, X., Paterson, A.H., 2013. PGDD: a database of gene and genome duplication in plants. Nucleic Acids Res. 41, D1152–8. doi: 10.1093/nar/gks1104

Lindahl, R., 1992. Aldehyde dehydrogenases and their role in carcinogenesis. Crit. Rev. Biochem. Mol. Biol. 27, 283–335. doi: 10.3109/10409239209082565

Liu, Z.-J., Sun, Y.-J., Rose, J., Chung, Y.-J., Hsiao, C.-D., Chang, W.-R., Kuo, I., Perozich, J., Lindahl, R., Hempel, J., Wang, B.-C., 1997. The first structure of an aldehyde dehydrogenase reveals novel interactions between NAD and the Rossmann fold. Nat. Struct. Mol. Biol. 4, 317–326. doi: 10.1038/nsb0497-317

Li, H., Rodda, M., Gnanasambandam, A., Aftab, M., Redden, R., Hobson, K., Rosewarne, G., Materne, M., Kaur, S., Slater, A.T., 2015. Breeding for biotic stress resistance in chickpea: progress and prospects. Euphytica 204, 257–288. doi: 10.1007/s10681-015-1462-8

Li, X., Guo, R., Li, J., Singer, S.D., Zhang, Y., Yin, X., Zheng, Y., Fan, C., Wang, X., 2013. Genome-wide identification and analysis of the aldehyde dehydrogenase (ALDH) gene superfamily in apple (*Malus × domestica* Borkh.). Plant Physiol. Biochem. 71, 268–282. doi: 10.1016/j.plaphy.2013.07.017

Lloyd, M.D., Boardman, K.D.E., Smith, A., van den Brink, D.M., Wanders, R.J.A., Threadgill, M.D., 2007. Characterisation of recombinant human fatty aldehyde dehydrogenase: implications for Sjögren-Larsson syndrome. J. Enzyme Inhib. Med. Chem. 22, 584–590. doi: 10.1080/14756360701425360

Matsuda, S., Vert, J.-P., Saigo, H., Ueda, N., Toh, H., Akutsu, T., 2005. A novel representation of protein sequences for prediction of subcellular location using support vector machines. Protein Sci. 14, 2804–2813. doi: 10.1110/ps.051597405

Millán, T., Madrid, E., Cubero, J.I., Amri, M., Castro, P., Rubio, J., 2015. Chickpea, in: De Ron, A.M. (Ed.), Grain Legumes, Handbook of Plant Breeding. Springer New York, New York, NY, pp. 85–109. doi: 10.1007/978-1-4939-2797-5_3

Missihoun, T.D., Kotchoni, S.O., 2018. Aldehyde dehydrogenases and the hypothesis of a glycolaldehyde shunt pathway of photorespiration. Plant Signal. Behav. 13, e1449544. doi: 10.1080/15592324.2018.1449544

Missihoun, T.D., Schmitz, J., Klug, R., Kirch, H.-H., Bartels, D., 2011. Betaine aldehyde dehydrogenase genes from *Arabidopsis* with different sub-cellular localization affect stress responses. Planta 233, 369–382. doi: 10.1007/s00425-010-1297-4

Parween, S., Nawaz, K., Roy, R., Pole, A.K., Venkata Suresh, B., Misra, G., Jain, M., Yadav, G., Parida, S.K., Tyagi, A.K., Bhatia, S., Chattopadhyay, D., 2015. An advanced draft genome assembly of a desi type chickpea (*Cicer arietinum* L.). Sci. Rep. 5, 12806. doi: 10.1038/srep12806

Perez-Miller, S.J., Hurley, T.D., 2003. Coenzyme isomerization is integral to catalysis in aldehyde dehydrogenase. Biochemistry 42, 7100–7109. doi: 10.1021/bi034182w

Plaxton, W.C., 1996. The organization and regulation of plant glycolysis. Annu. Rev. Plant Physiol. Plant Mol. Biol. 47, 185–214. doi: 10.1146/annurev.arplant.47.1.185

Prochnik, S.E., Umen, J., Nedelcu, A.M., Hallmann, A., Miller, S.M., Nishii, I., Ferris, P., Kuo, A., Mitros, T., Fritz-Laylin, L.K., Hellsten, U., Chapman, J., Simakov, O., Rensing, S.A., Terry, A., Pangilinan, J., Kapitonov, V., Jurka, J., Salamov, A., Shapiro, H., Rokhsar, D.S., 2010. Genomic analysis of organismal complexity in the multicellular green alga *Volvox carteri*. Science 329, 223–226. doi: 10.1126/science.1188800

Rejeb, I.B., Pastor, V., Mauch-Mani, B., 2014. Plant responses to simultaneous biotic and abiotic stress: molecular mechanisms. Plants 3, 458–475. doi: 10.3390/plants3040458

Richardson, J.S., Arendall, W.B., Richardson, D.C., 2003. New Tools and Data for Improving Structures, Using All-Atom Contacts, in: Macromolecular Crystallography, Part D, Methods in Enzymology. Elsevier, pp. 385–412. doi: 10.1016/S0076-6879(03)74018-X

Rodrigues, S.M., Andrade, M.O., Gomes, A.P.S., Damatta, F.M., Baracat-Pereira, M.C., Fontes, E.P.B., 2006. *Arabidopsis* and tobacco plants ectopically expressing the soybean antiquitin-like ALDH7 gene display enhanced tolerance to drought, salinity, and oxidative stress. J. Exp. Bot. 57, 1909–1918. doi: 10.1093/jxb/erj132

Shin, J.-H., Kim, S.-R., An, G., 2009. Rice aldehyde dehydrogenase7 is needed for seed maturation and viability. Plant Physiol. 149, 905–915. doi: 10.1104/pp.108.130716

Skibbe, D.S., Liu, F., Wen, T.-J., Yandeau, M.D., Cui, X., Cao, J., Simmons, C.R., Schnable, P.S., 2002. Characterization of the aldehyde dehydrogenase gene families of *Zea mays* and *Arabidopsis*. Plant Molecular Biology 48, 751–764.

Stiti, N., Missihoun, T.D., Kotchoni, S.O., Kirch, H.-H., Bartels, D., 2011. Aldehyde Dehydrogenases in *Arabidopsis thaliana*: Biochemical Requirements, Metabolic Pathways, and Functional Analysis. Front. Plant Sci. 2, 65. doi: 10.3389/fpls.2011.00065

Tamura, K., Stecher, G., Peterson, D., Filipski, A., Kumar, S., 2013. MEGA6: Molecular Evolutionary Genetics Analysis version 6.0. Mol. Biol. Evol. 30, 2725–2729. doi: 10.1093/molbev/mst197

Tan, S., Wu, S., 2012. Genome Wide Analysis of Nucleotide-Binding Site Disease Resistance Genes in *Brachypodium distachyon*. Comp. Funct. Genomics 2012, 418208. doi: 10.1155/2012/418208

Tian, F.-X., Zang, J.-L., Wang, T., Xie, Y.-L., Zhang, J., Hu, J.-J., 2015. Aldehyde Dehydrogenase Gene Superfamily in *Populus*: Organization and Expression Divergence between Paralogous Gene Pairs. PLoS ONE 10, e0124669. doi: 10.1371/journal.pone.0124669

Varshney, R.K., Song, C., Saxena, R.K., Azam, S., Yu, S., Sharpe, A.G., Cannon, S., Baek, J., Rosen, B.D., Tar’an, B., Millan, T., Zhang, X., Ramsay, L.D., Iwata, A., Wang, Y., Nelson, W., Farmer, A.D., Gaur, P.M., Soderlund, C., Penmetsa, R.V., Cook, D.R., 2013. Draft genome sequence of chickpea (*Cicer arietinum*) provides a resource for trait improvement. Nat. Biotechnol. 31, 240–246. doi: 10.1038/nbt.2491

Vasiliou, V., Bairoch, A., Tipton, K.F., Nebert, D.W., 1999. Eukaryotic aldehyde dehydrogenase (ALDH) genes: human polymorphisms, and recommended nomenclature based on divergent evolution and chromosomal mapping. Pharmacogenetics 9, 421–434.

Wang, L., Guo, Z., Zhang, Y., Wang, Y., Yang, G., Yang, L., Wang, Li, Wang, R., Xie, Z., 2017. Overexpression of LhSorNPR1, a NPR1-like gene from the oriental hybrid lily “Sorbonne”, conferred enhanced resistance to *Pseudomonas syringae* pv. tomato DC3000 in Arabidopsis. Physiol. Mol. Biol. Plants 23, 793–808. doi: 10.1007/s12298-017-0466-3

Wang, W., Jiang, W., Liu, J., Li, Yang, Gai, J., Li, Yan, 2017. Genome-wide characterization of the aldehyde dehydrogenase gene superfamily in soybean and its potential role in drought stress response. BMC Genomics 18, 518. doi: 10.1186/s12864-017-3908-y

Wood, A.J., Duff, R.J., 2009. The aldehyde dehydrogenase (ALDH) gene superfamily of the moss *Physcomitrella patens* and the algae *Chlamydomonas reinhardtii* and *Ostreococcus tauri*. Bryologist 112, 1–11. doi: 10.1639/0007-2745-112.1.1

Xu, X., Guo, R., Cheng, C., Zhang, H., Zhang, Y., Wang, X., 2013. Overexpression of ALDH2B8, an aldehyde dehydrogenase gene from grapevine, sustains *Arabidopsis* growth upon salt stress and protects plants against oxidative stress. Plant Cell Tiss. Organ Cult. 114, 187–196. doi: 10.1007/s11240-013-0314-2

Yoshiba, Y., Kiyosue, T., Nakashima, K., Yamaguchi-Shinozaki, K., Shinozaki, K., 1997. Regulation of levels of proline as an osmolyte in plants under water stress. Plant Cell Physiol. 38, 1095–1102.

Yoshida, A., Rzhetsky, A., Hsu, L.C., Chang, C., 1998. Human aldehyde dehydrogenase gene family. Eur. J. Biochem. 251, 549–557.

Zahniser, M.P.D., Prasad, S., Kneen, M.M., Kreinbring, C.A., Petsko, G.A., Ringe, D., McLeish, M.J., 2017. Structure and mechanism of benzaldehyde dehydrogenase from *Pseudomonas putida* ATCC 12633, a member of the Class 3 aldehyde dehydrogenase superfamily. Protein Eng. Des. Sel. 30, 271–278. doi: 10.1093/protein/gzx015

Zhang, C.-R., Huang, X.-L., Wu, J.-Y., Feng, B.-H., Chen, Y.-F., 2006. Identification of thidiazuron-induced ESTs expressed differentially during callus differentiation of alfalfa (*Medicago sativa*). Physiol. Plant. 128, 732–739. doi: 10.1111/j.1399-3054.2006.00763.x

Zhang, N., Zoltner, M., Leung, K.-F., Scullion, P., Hutchinson, S., Del Pino, R.C., Vincent, I.M., Zhang, Y.-K., Freund, Y.R., Alley, M.R.K., Jacobs, R.T., Read, K.D., Barrett, M.P., Horn, D., Field, M.C., 2018. Host-parasite co-metabolic activation of antitrypanosomal aminomethyl-benzoxaboroles. PLoS Pathog. 14, e1006850. doi: 10.1371/journal.ppat.1006850

Zhang, Y., Mao, L., Wang, H., Brocker, C., Yin, X., Vasiliou, V., Fei, Z., Wang, X., 2012. Genome-wide identification and analysis of grape aldehyde dehydrogenase (ALDH) gene superfamily. PLoS ONE 7, e32153. doi: 10.1371/journal.pone.0032153

